# RhoGAP19D inhibits Cdc42 laterally to control epithelial cell shape and prevent invasion

**DOI:** 10.1101/2020.09.17.300145

**Authors:** Weronika Fic, Rebecca Bastock, Francesco Raimondi, Erinn Los, Yoshiko Inoue, Jennifer Gallop, Robert B. Russell, Daniel St Johnston

## Abstract

Cdc42-GTP is required for apical domain formation in epithelial cells where it recruits and activates the Par-6/aPKC polarity complex, but how the activity of Cdc42 itself is restricted apically unclear. We used sequence analysis and 3D structure modelling to determine which *Drosophila* GTPase Activating Proteins (GAPs) are likely to interact with Cdc42 and identified RhoGAP19D as the sole Cdc42GAP required for polarity in the follicle cell epithelium. RhoGAP19D is recruited by α-catenin to lateral E-cadherin adhesion complexes, resulting in exclusion of active Cdc42 from the lateral domain. *rhogap19d* mutants therefore lead to lateral Cdc42 activity, which expands the apical domain through increased Par-6/aPKC activity and stimulates lateral contractility through the myosin light chain kinase, Genghis khan (MRCK). This causes buckling of the epithelium and invasion into the adjacent tissue, a phenotype resembling that of pre-cancerous breast lesions. Thus, RhoGAP19D couples lateral Cadherin adhesion to the apical localisation of active Cdc42, thereby suppressing epithelial invasion.

## Introduction

The form and function of epithelial cells depends on their polarisation into distinct apical, lateral and basal domains by conserved polarity factors (Rodriguez-Boulan and Macara, 2014; St Johnston and Ahringer, 2010). This polarity is then maintained by mutual antagonism between apical polarity factors like atypical protein kinase C (aPKC) and lateral factors, such as Lgl and Par-1. While many aspects of the polarity machinery are now well understood, it is still unclear how the apical domain is initiated and what role Cdc42 plays in this process.

Cdc42 was identified for its role in establishing polarity in budding yeast, where it targets cell growth to the bud tip by polarising the actin cytoskeleton and exocytosis towards a single site (Chiou et al., 2017). It has subsequently been found to function in the establishment of cell polarity in multiple contexts. For example, Cdc42 recruits and activates the anterior PAR complex to polarise the anterior-posterior axis in the *C. elegans* zygote and the apical-basal axis during the asymmetric divisions of *Drosophila* neural stem cells (Gotta et al., 2001; Kay and Hunter, 2001; Atwood et al., 2007; Rodriguez et al., 2017).

Cdc42 also plays an essential role in the apical-basal polarisation of epithelial cells, where it is required for apical domain formation (Genova et al., 2000; Hutterer et al., 2004; Jaffe et al., 2008; Bray et al., 2011; Fletcher et al., 2012). Cdc42 is active when bound to GTP, which changes its conformation to allow it to bind downstream effector proteins that control the cytoskeleton and membrane trafficking. An important Cdc42 effector in epithelial cells is the Par-6/aPKC complex. Par-6 binds directly to the switch 1 region of Cdc42 GTP through its semi-CRIB domain (Cdc42 and Rac interactive binding) (Lin et al., 2000; Joberty et al., 2000; Qiu et al., 2000; Yamanaka et al., 2001). This induces a change in the conformation of Par-6 that allows it to bind to the C-terminus of another key apical polarity factor, the transmembrane protein, Crumbs, which triggers the activation of aPKC’s kinase activity (Dong et al., 2019; Peterson et al., 2004; Whitney et al., 2016). As a result, active aPKC is anchored to the apical membrane, where it phosphorylates and excludes lateral factors, such as Lgl, Par-1 and Bazooka (Betschinger et al., 2003; Hurov et al., 2004; Suzuki et al., 2004; Nagai-Tamai et al., 2002; Morais-de-Sá et al., 2010). In addition to this direct role in apical-basal polarity, Cdc42 also regulates the organisation and activity of the apical cytoskeleton through effectors such as N-WASP, which promotes actin polymerisation, and MRCK (*Gek* in *Drosophila*), which phosphorylates the myosin regulatory light chain to activate contractility (Padrick and Rosen, 2010; Zihni et al., 2017).

This crucial role of active Cdc42 in specifying the apical domain raises the question of how Cdc42-GTP itself is localised apically. In principle, this could involve activation by Cdc42GEFs that are themselves apical or lateral inactivation by Cdc42GAPs. The Cdc42GEFs, Tuba, Intersectin 2 and Dbl3 have been implicated in activating Cdc42 in mammalian epithelia (Oda et al., 2014; Otani et al., 2006; Qin et al., 2010; Rodriguez-Fraticelli et al., 2010; Zihni et al., 2014). Only Dbl3 localises apical to tight junctions, however, as Tuba is cytoplasmic and enriched at tricellular junctions and Intersectin 2 localises to centrosomes. Thus, GEF activity may not be exclusively apical, suggesting that it is more important to inhibit Cdc42 laterally. Although nothing is known about the role of GAPs in restricting Cdc42 activity to the apical domain of epithelial cells, this mechanism plays an instructive role in establishing radial polarity in the blastomeres of the early *C. elegans* embryo. In this system, the Cdc42 GAP, PAC-1, is recruited by the Cadherin adhesion complex to sites of cell-cell contact, thereby restricting active Cdc42 and its effector the Par-6/aPKC complex to the contact-free surface (Anderson et al., 2008; Klompstra et al., 2015).

Here we analysed the roles of Cdc42GAPs in epithelial polarity using the follicle cells that surround developing *Drosophila* egg chambers as a model system (Bastock and St Johnston, 2008). By generating mutants in a number of candidate Cdc42 GAPs, we identified the Pac-1 orthologue, RhoGAP19D, as the GAP that restricts active Cdc42 to the apical domain. In the absence of RhoGAP19D, lateral Cdc42 activity leads to an expansion of the apical domain and a high frequency of epithelial invasion into the germline tissue, a phenotype that mimics the early steps of carcinoma formation.

## Results

To confirm that Cdc42 regulates apical domain formation in *Drosophila* epithelia, we generated homozygous mutant clones of *cdc42*^2^, a null allele, in the follicle cell epithelium that surrounds developing egg chambers (Fig 1A). Mutant cells lose their cuboidal shape, leading to gaps and multi-layering in the epithelium, and fail to localise GFP-aPKC apically, indicating that Cdc42 is required for polarity in the follicle cell layer.

**Figure 1.**
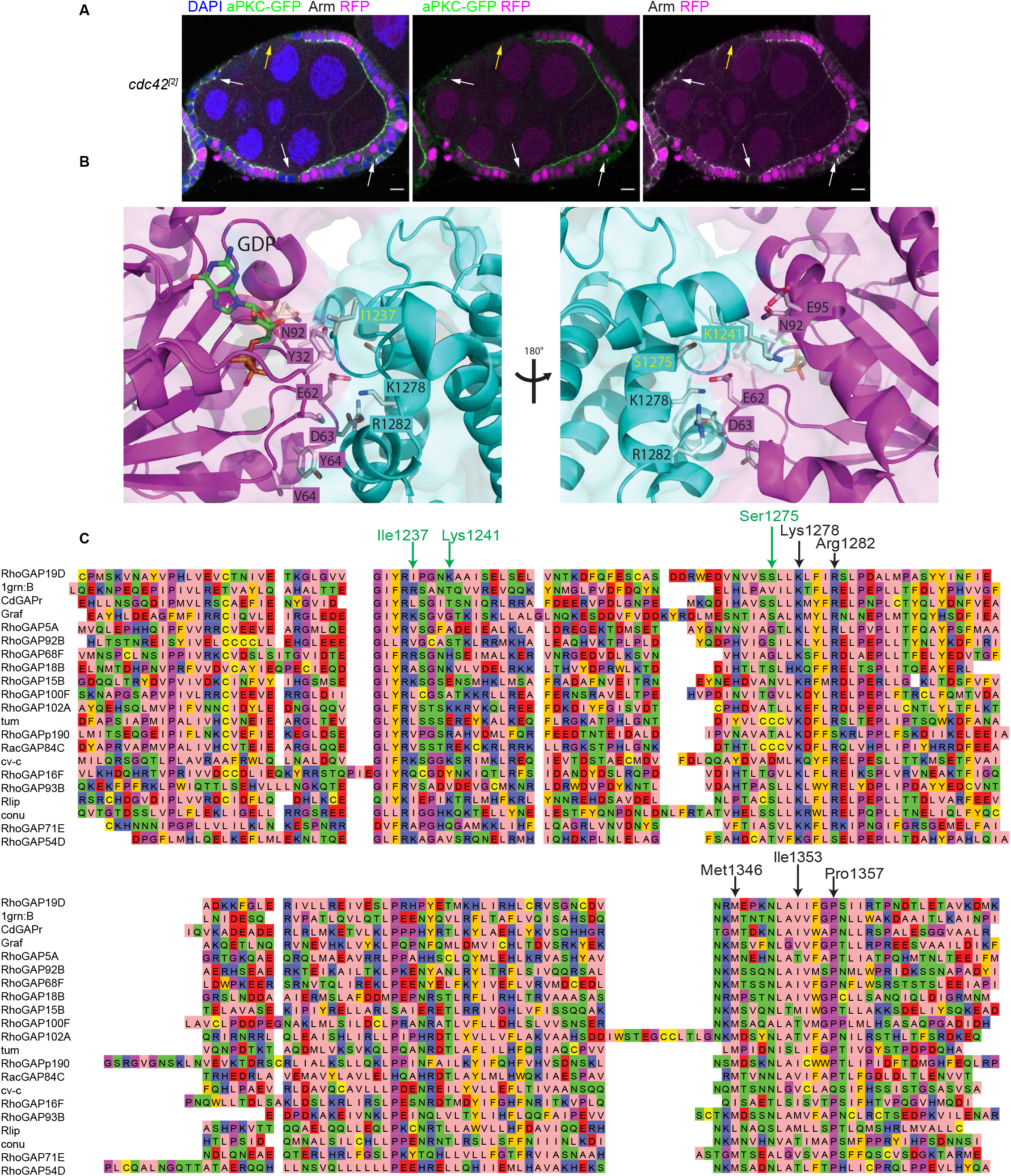
Cdc42 is essential for the establishment of cell polarity in Drosophila follicle cells. A) stage 7 egg chamber containing multiple *cdc42*^[2]^ mutant follicle cell clones (marked by the loss of RFP, magenta) stained for Armadillo (white) and DAPI (blue) expressing endogenously-tagged GFP-aPKC (green). The mutant cells are marked by arrows. GFP-aPKC is lost from the apical side of c*dc42* mutant follicle cells. In some cells, GFP-aPKC co-localises with Armadillo in puncta. Cells lacking Cdc42 become round and often lose contact with neighbouring cells, resulting in breaks in the epithelial layer. In other cases, the *cdc42* mutant cells lie basally to to wild-type cells. Scale bars 10μm. B) A diagram showing the interface between Cdc42 and bound ARHGAP1/RhoGAP19D. Key amino acids that mediate the interaction are shown in purple for Cdc42 and in green for ARHGAP1. Amino acid changes that are predicted to strengthen the interaction between Cdc42 and RhoGAP19D are shown in parentheses. C) Alignment of Drosophila GAPs and human ARHGAP1. Conserved amino acids involved in the interaction with Cdc42 are indicated by black arrows. The green arrows mark the variable amino acids that are predicted to strengthen the interaction between Cdc42 and RhoGAP19D.

There are 22 Rho GTPase activating proteins in the *Drosophila* genome (Table S1), but in most cases, it is unclear whether they regulate Rho, Rac or Cdc42. We therefore predicted the tendency for each of the Drosophila GAPs to interact with Cdc42 using InterPReTS (Aloy and Russell, 2002). This uses a known structure of a protein complex (in this case the structure of the human CDC42/ARHGAP1 complex, (Nassar et al., 1998) as a template to predict whether homologous proteins (in this case other *Drosophila* GAPs and Cdc42) would be able to interact in the same way. The fit of each sequence pair on the structure is assessed via statistical potentials that score the compatibility of each amino acid pair at the (e.g. GAP/Cdc42) interface. The ranks for each *Drosophila* GAP according to its likelihood of interacting with *Drosophila* Cdc42 is shown in Table S2 Interestingly, the *Drosophila* ARHGAP1 orthologue, RhoGAP68F was only second in the ranking behind another known Cdc42GAP, the PAC-1orthologue, RhoGAP19D. Inspection of the InterPreTS results in detail shows that several key, conserved positions mediating interactions in the structure of the human CDC42/ARHGAP1 complex are conserved in RhoGAP19D (and many other) *Drosophila* GAPs, in addition to three positions that appear to make the interaction stronger (Figure 1). Specifically, an Arg in ARHGAP1 is replaced by a Ile (1237 in RhoGAP19D) making for a better interaction with Ala13 in Cdc42, a Val is replaced by a Ser (1275) making a more favourable interaction with Glu62, and a Thr is replaced by a Lys (1237) possibly making an additional salt bridge with Glu95 and additional interactions with Asn92.

To test whether any of these putative Cdc42 GAPs play a role in epithelial polarity, we generated null mutants in the seven highest ranked GAPs using CRISPR-mediated mutagenesis (Table S3) and examined their phenotypes in the follicle cell epithelium that surrounds developing egg chambers. Mutants in RhoGAP92B are lethal and we therefore used the Flp/FRT system to generate homozygous mutant follicle cell clones, whereas mutants in the other GAPs are homozygous viable or semi-viable, allowing us to analyse the follicle cell phenotype in homozygous mutant females. Null mutants in *RhoGAP68F*, *CdGAPr*, *RhoGAP92B*, *RhoGAP82C*, *conundrum*(Neisch et al., 2013) and *RhoGAP93B* cause no discernible changes in follicle cell shape or polarity, as shown by the localisation of aPKC apically and Lgl laterally (Fig S2). By contrast, 40% of homozygous mutant *RhoGAP19D* egg chambers show invasions of regions of the follicular epithelium into the overlying germline cyst (Figure 2A-C). This invasive phenotype is not a consequence of over-proliferation of the mutant follicle cells, because mutant egg chambers contain the same number of follicle cells as wild-type egg chambers and the same proportion of homozygous mutant cells and wild-type cells are in mitosis at stages 4-5 (Figure 2 D-F). Introducing endogenously tagged E-cadherin into the *RhoGap19D* mutant background reveals that the invading follicle cells maintain their apical adherens junctions, indicating that they have not undergone an epithelial to mesenchymal transition and are still epithelial in nature (Figure 2G).

**Figure 2.**
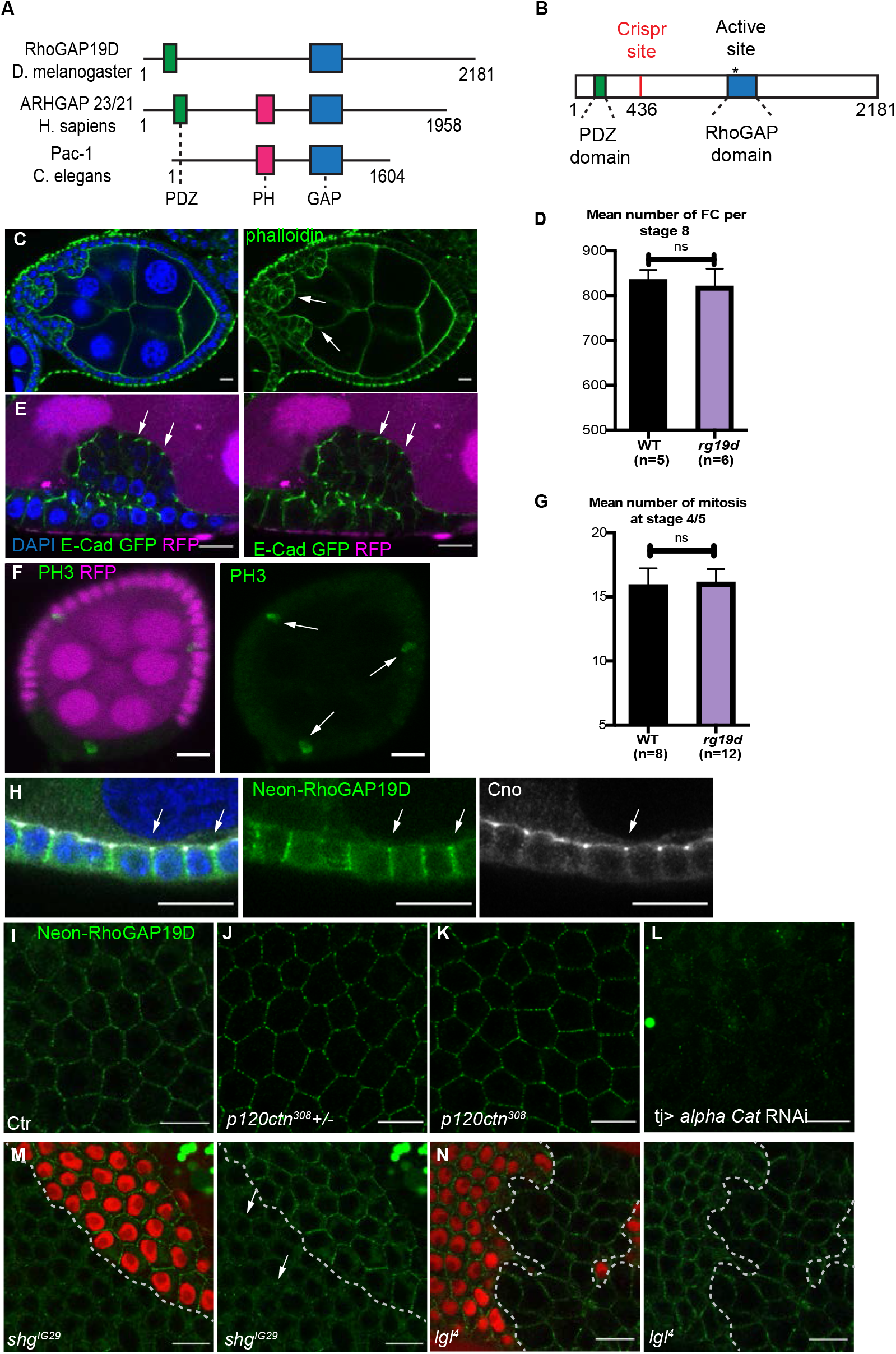
RhoGAP19D is required for the integrity of epithelial layer. A) Comparison of the domain structure of *Drosophila* RhoGAP19D with its orthologues, human ARHGAP23/21 and *C. elegans* PAC-1. RHDOGAP19D contains PDZ and GAP domains, but lacks a PH domain. B) Diagram showing the CRISPR-induced mutations in RhoGAP19D. The mutations generate proteins that lack the GAP domain, but still contain the PDZ domain. C) A stage 7 *rhogp19d* mutant egg chamber showing invasions of the follicle cell epithelium into the adjacent germ line. The mutant cells (white arrows) invade between the nurse cells at the anterior of the egg chamber. Phalloidin staining is shown in green and DAPI in blue. D) Graph showing the number of wild-type and *rhogap19d* mutant cells per egg chamber at stage 8. E) Invading *rhogap19d* mutant cells (marked by the loss of RFP, magenta) maintain their adherens junctions and epithelial organisation, as shown by E-cadherin-GFP expression; DAPI (blue). F) PH3 staining (green) of mitotic cells in a stage 5 egg chamber containing a *rhogap19d* mutant clone (marked by the loss of RFP, magenta). There is no increase in cell division in the mutant clone. G) Quantification of the number of mitoses in wild-type and *rhogap19d* mutant egg chambers during stages 4-5. H) mNeonGreen-RHOGAP19D (green)localises to the lateral domain of the follicle cells and is slightly enriched at the adherens junctions stained with Cno (white); DAPI (blue). RHOGAP19D protein in the germline was depleted by RNAi. I) Surface view of wild-type cells expressing mNeonGreen-RhoGAP19D J) and K) mNeonGreen-RHOGAP19D localises normally in *p120 catenin*^308^/+ and *p120 catenin*^308^ homozygous cells. L and M) mNeonGreen-RHOGAP19D recruitment to lateral domain is almost lost in cells expressing α-catenin RNAi (L) and is strongly reduced in *shotgun*^IG29^ mutant clones (marked by the loss of RFP, magenta, M). N) mNeonGreen-RHOGAP19D localisation is not affected in *lgl*^4^ mutant clones (marked by the loss of RFP, magenta) compared to wild-type cells. Scale bars 10μm.

We examined the localisation of RhoGAP19D protein by using CRISPR-mediated homologous recombination to insert the mNeonGreen fluorescent tag at the N-terminus of the endogenous RhoGAP19D coding region. Neon::RhoGAP19D localises laterally in the follicle cells, covering the full length of the domain including the apical adherens junctions, where it sometimes appears to be slightly enriched (Figure 2h and i). A similar lateral localisation is observed in the salivary gland and the testis accessory gland, indicating that RhoGAP19D is a lateral factor in multiple epithelia (Figure S3).

The *C. elegans* orthologue of RhoGAP19D, PAC-1, is recruited to cell contacts by the E-Cadherin complex through redundant interactions with α-catenin and p120-catenin (Klompstra et al., 2015). We observed no change in the lateral recruitment of RhoGAP19D in *p120 catenin* null mutants, but the junctional signal was almost completely lost when α-catenin was depleted by RNAi (Figure 2i-l, Figure SXa). Thus, RhoGAP19D is localised to the lateral membrane by a non-redundant interaction with α-catenin, which links it to Cadherin adhesion complexes. The lateral localisation of RhoGAP19D was strongly reduced in clones homozygous for *shotgun*^IG29^, a null mutant in *shotgun* (E-cadherin), whereas clones homozygous for a deletion of N-cadherin 1 and N-cadherin 2 had no effect(Tepass et al., 1996; Prakash et al., 2005). The weaker phenotype of *shotgun*^IG29^ clones compared to α-catenin knockdown is presumably because N-cadherin is up-regulated in E-cadherin mutants and either E- or N-Cadherin can recruit α-catenin and RhoGAP19D (Grammont, 2007). α-catenin and E-cadherin are concentrated in the apical adherens junctions, whereas RhoGAP19D shows only a slight apical enrichment and is much more uniformly distributed along the lateral membrane. Since this suggests that other factors may modulate the recruitment of RhoGAP19, we examined whether any lateral polarity factors affect its distribution, but observed no change when the lateral adhesion proteins, FasII, FasIII or Neuroglian are knocked down by RNAi or in null mutant clones for the lateral polarity factor, Lgl (Fig 2m, Fig S2b-g).

The localisation of RhoGAP19D suggests that it may function to inhibit Cdc42 laterally. We therefore examined where Cdc42 is active by following the localisation of an endogenously tagged version of the Cdc42 effector, N-WASP (Kim et al., 2000). In wild-type cells, N-WASP-Neon localises exclusively to the apical domain, consistent with the apical localisation of active Cdc42. By contrast, N-WASP also localizes along the lateral membrane in *rhogap19d* mutant clones (Fig 3a,b). Thus, RhoGAP19D functions to inhibit Cdc42 laterally and is the only GAP that fulfills this role in the follicle cells.

**Figure 3.**
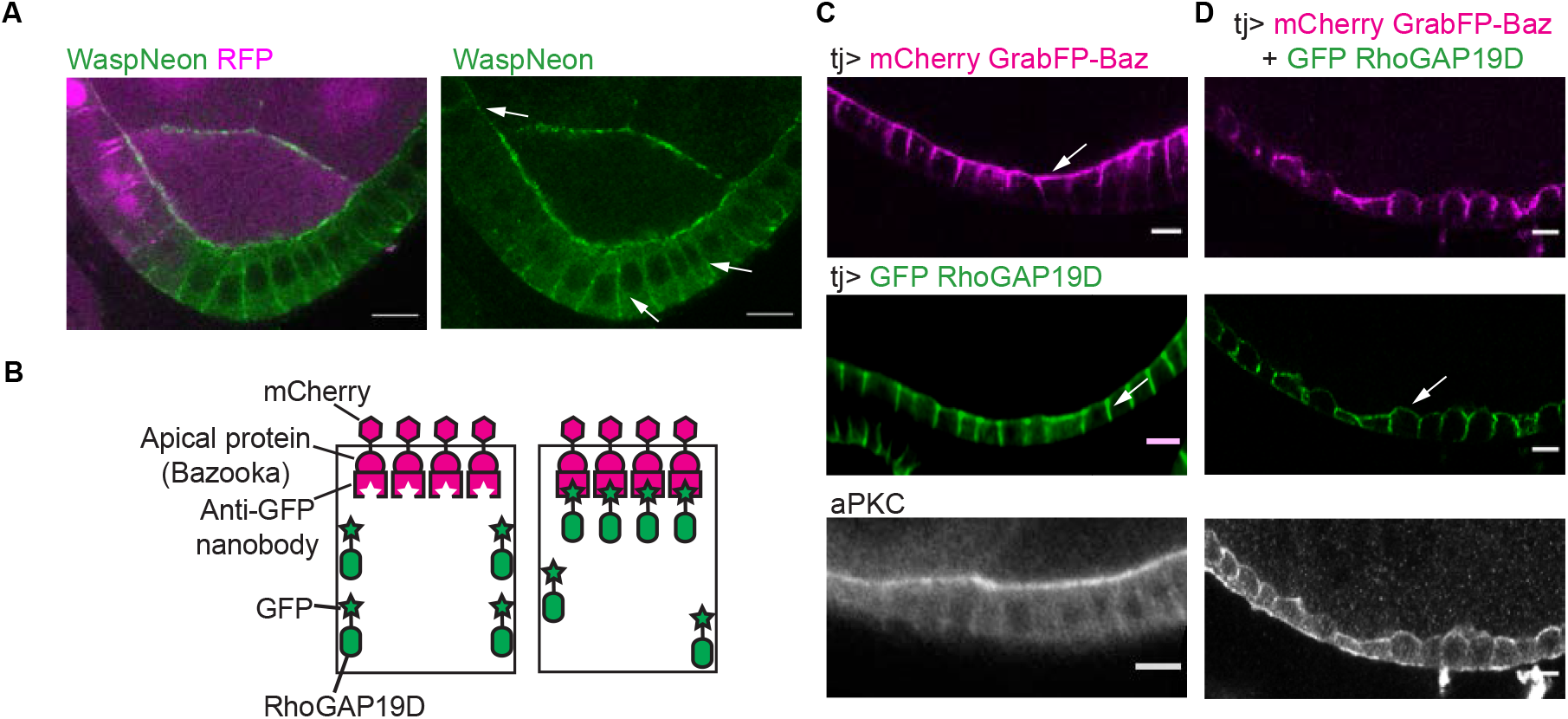
RHOGAP19D inhibits CDC42 activity laterally. A) The CDC42 effector, N-WASP (tagged with mNeonGreen) spreads laterally in *rhogap19D* mutant cells (marked by the loss of RFP, magenta). B) Cartoon showing the UAS-GrabFP-A-int system. C) mCherry-Grab FP Bazooka predominantly localizes to apical side of the follicle cells when expressed under the control of TJ-Gal4, whereas GFP-RhoGAP19D alone localizes laterally. D) Co-expression of mCherry-Grab FP Baz and GFP-RhoGAP19D results in the apical recruitment of RhoGAP19D, leading to a loss of epithelial and mis-localisation of aPKC (white). Scale bars 10μm.

To confirm that RhoGAP19D represses Cdc42 activity, we used UAS-GrabFP-A-int to mislocalise the protein to the apical domain (Harmansa et al., 2017). The GrabFP-A-int construct consists of an N-terminal Cherry, a transmembrane domain and an anti-GFP nanobody fused to the localisation signal of Bazooka (Par-3). When this construct is expressed in the follicle cells under the control of Tj-Gal4, the fusion protein localises to the apical membrane and apical junctions without any apparent effect on the appearance of the cells (Fig 3A). Similarly, over-expression of UAS-GFP-RhoGAP19D alone results in higher levels of RhoGAP19D along the lateral membrane, but has no effect on cell polarity or morphology during stages 1-8 of oogenesis. When GFP-RhoGAP19D and GrabFP-A-int are co-expressed, however, the apical recruitment of GFP-RhoGAP19D by the anti-GFP nanobody disrupts polarity and epithelial organization, as shown by the failure to concentrate aPKC apically and the irregular cell shapes (Fig 3E,F). This phenotype closely resembles that of *cdc42* mutants, providing further evidence RhoGAP19D is a specific Cdc42 GAP.

To investigate the cellular basis for the invasive behaviour of *rhogap19d* mutant follicle cells, we compared the phenotypes of mutant and wildtype cells in the same epithelium by generating homozygous mutant clones. Live imaging reveals that *rhogap19d* mutant cells are taller than wild-type cells, with dome-shaped apical surfaces that protrude into the germ line (Figure 4A). The distance between the apical and basal surfaces of mutant cells is 20% greater than wild-type in fixed samples (Figure 4b,e). However, the adherens junctions, marked by E cadherin-GFP and Canoe, do not show a corresponding change in position, and remain level with or slightly below the adherens junctions in the neighbouring wild-type cells (Figure 4b,c, g). The adherens junctions mark the boundary between the apical and lateral domains, suggesting that the apical domain has expanded. This is indeed the case, as GFP-aPKC localises all over the domed region of membrane above the adherens junctions, which is 40% longer than in wild-type. (Figure 4c, f). The apical transmembrane protein Crumbs show a similar extension across the expanded apical domain, but is more enriched in the sub-apical region above the adherens junctions, reflecting its accumulation in regions where it can engage in homophilic interactions with Crumbs in adjacent cells (Thompson et al., 2013) (Figure 4c). By contrast, the lateral domain, marked by Lgl-GFP, is decreased in length (Figure 4d, g). Lateral Cdc42 activity therefore expands the apical domain at the expense of the lateral domain to generate taller cells that protrude into the germ line. A similar apical expansion is also observed in *rhogap19d* mutant testis accessory glands, suggesting that this is a general phenotype of loss of RhoGAP19D in *Drosophila* epithelia (Fig S3).

**Figure 4.**
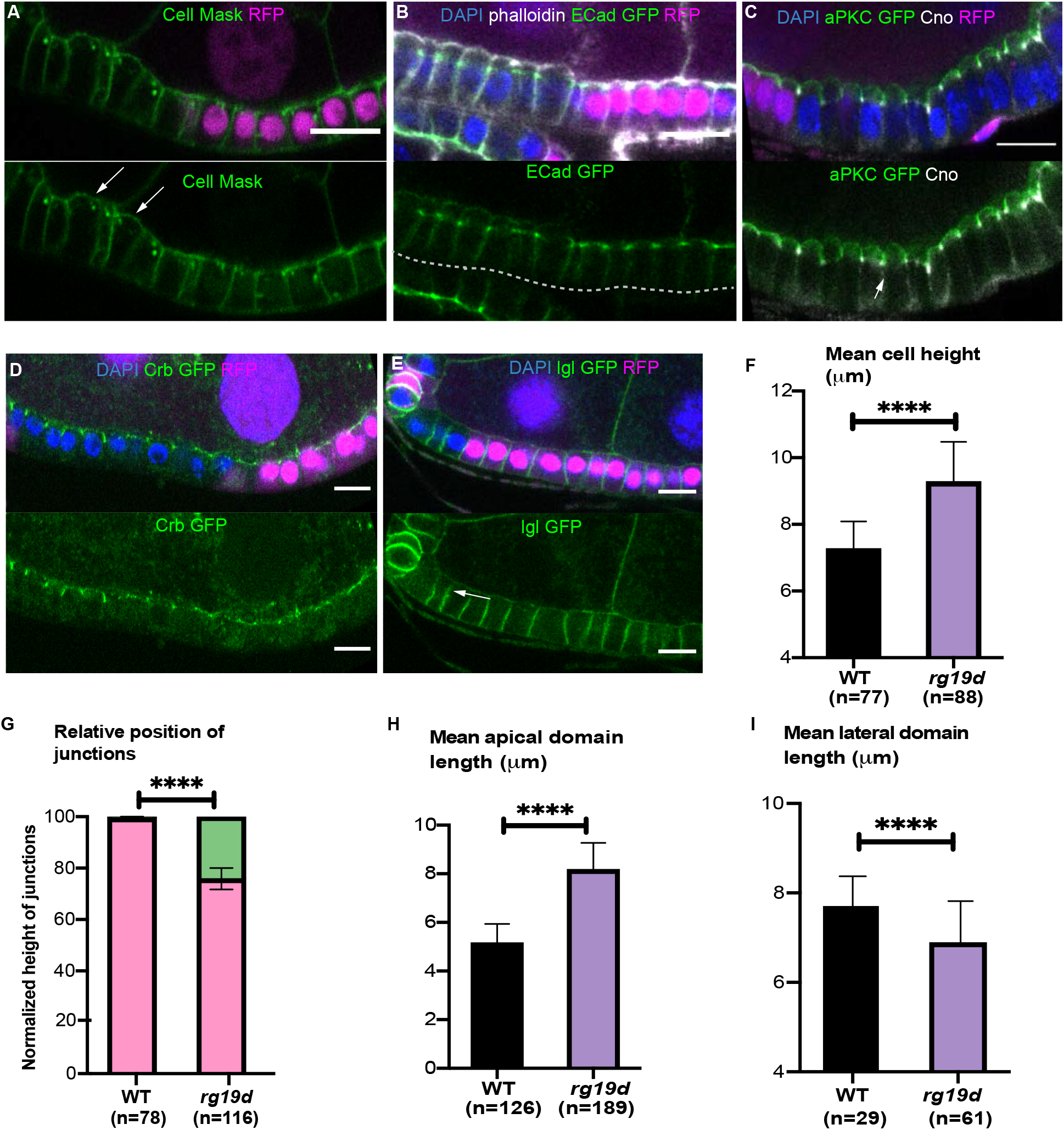
*rhogap19D* mutant cells are taller and have an enlarged apical domain. A-E) Regions of the stage 7 follicle cell epithelium stained with DAPI (blue) containing clones of *rhogap19d* mutant cells marked by the loss of RFP (magenta). (A) Cell mask staining (green) of the plasma membranes reveals that mutant cells are taller than wild-type cells and have domed apical surfaces. The arrows indicate the apical surfaces of the mutant cells. B) The adherens junctions marked by endogenously-tagged E-Cad-GFP (green) form at the same level in *rhogap19d* mutant and wild-type cells. (Phalloidin; white). C) In *rhogap19d* mutant cells, GFP-aPKC localises all around the apical domain above the adherens junctions (marked by Cno staining; white). D) Crb-GFP marks an enlarged sub-apical region in *rhogap19d* cells. E) *rhogap19d* mutant cells have slightly shorter lateral domains than wild-type cells, as shown by Lgl-GFP localisation (green). F) A graph showing the mean cell height in wild-type and *rhogap19d* mutant cells. G) A graph showing the relative position of adherens junctions compared to total cell height in wild-type and *rhogap19d* mutant cells. H) A graph showing the mean apical domain length in wild-type and *rhogap19d* mutant cells. I) A graph showing the mean lateral domain length in wild-type and *rhogap19d* mutant cells. Scale bars 10μm.

To gain insight into how *rhogap19d* mutant follicle cells invade the germ line, we imaged living egg chambers at stages 5-7, the stages when invasions are most likely to occur (Movie S1, Figure 5a). Not only are the mutant cells taller than wild-type with domed apical surfaces, but they are also more motile. Temporal projections show that the mutant cells expand and contract along their apical-basal axes, whereas wild-type cells are static (Figure 5b). The apical expansion of the mutant cells and the up and down movements are likely to increase strain in the epithelium and raise the probability of regions of the follicle cell layer buckling towards the germ line (Figure 5a). More rarely, we observe clusters of cells that have detached from the basement membrane and are beginning to invade (Figure 5c).

**Figure 5.**
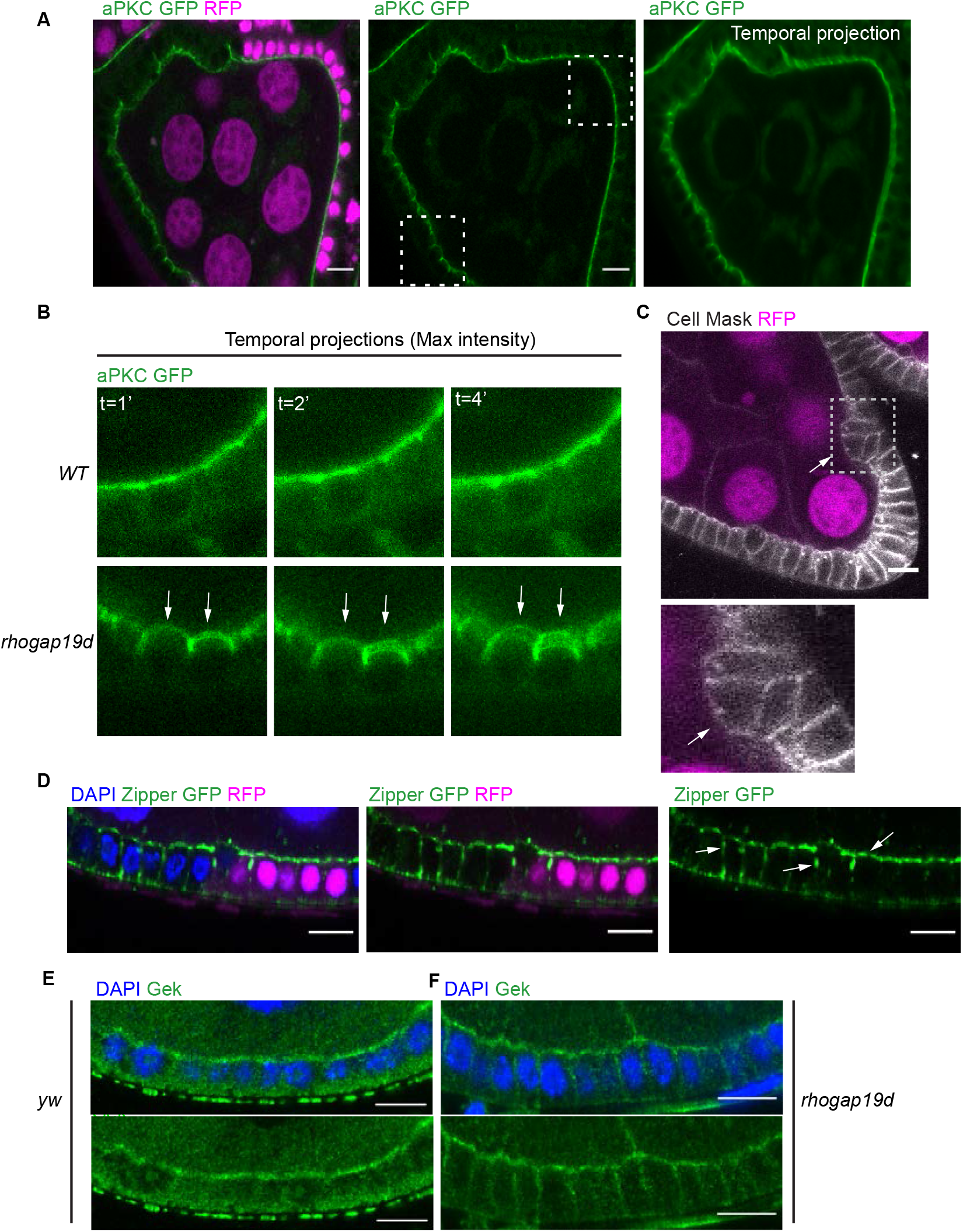
*rhogap19d* mutant cells are more motile and contract laterally. A) Single frames and a temporal projection of a time-lapse movie of a stage 7 egg chamber containing a large *rhogap19d* mutant clone (marked by the loss of RFP; magenta) and expressing GFP-aPKC (green). The blurred apical surfaces of the mutant cells in the temporal projection indicate that they are moving between frames B) Magnification of the boxed areas in (A), showing that *rhogap19d* mutant cells (bottom panels) are more motile than wild-type cells (top panels). Temporal projections after 1, 2 and 4 mins. C) A single frame from a movie showing a cluster of *rhogap19d* mutant cells (marked by the loss of RFP, magenta) beginning to invade the germ line. The cells in the cluster appear to have detached from the basement membrane. (Cell mask; white). The lower panel shows a magnification of the boxed area. D) *rhogap19d* mutant cells have lateral foci of non-muscle Myosin II foci (Zipper-GFP; green) and reduced levels at the apical side, compared to wild-type cells. (DAPI; blue). E) Gek (green) localises apically in wild-type follicle cells, but extends along the lateral domain of *rhogap19d* mutant cells. DAPI (blue). Scale bars 10μm.

The higher motility suggests that myosin activity is increased in mutant cells, and we therefore examined the distribution of non-muscle myosin II (NMYII) using a protein trap insertion in the heavy chain (Zipper). This revealed that the mutant cells have more numerous and larger NMYII foci along their lateral membranes and reduced levels of apical NMYII (Figure 5d). This increase in lateral NMYII is likely to account for the apical-basal contractions in mutant cells. In MDCK cells, Cdc42 recruits and activates NMYII apically through its effector, Myotonic dystrophy kinase-related Cdc42-binding kinase (MRCK), which phosphorylates the myosin regulatory light chain to stimulate contractility (Zihni et al., 2017; Zhao and Manser, 2015). This suggests that the *Drosophila* orthologue of MRCK, Genghis kan (Gek) might play a similar role in coupling Cdc42 to the activation of NMYII in the follicle cell epithelium. Antibody staining revealed that Gek is predominantly localised to the apical surface of the follicle cells, consistent with its role in MDCK cells (Figure 5e). In *rhogap19d* mutants, however, Gek extends along the lateral membrane (Figure 5f). Thus, the ectopic Cdc42 activity in *rhogap19d* mutants recruits Gek to the lateral membrane, where it can localise and activate NMYII.

Our results suggest that the invasive phenotype of *rhogap19d* mutants depends on a partial disruption of polarity, in which the apical domain expands at the expense of the lateral domain. Since the relative sizes of the apical and lateral domains are determined by mutual antagonism between apical and lateral polarity factors, reducing the dosage of lateral factors should enhance this phenotype, whereas reducing apical factors should suppress it. We therefore tested whether polarity mutants act as dominant modifiers of the *rhogap19d* phenotype (Figure 6). Removing one copy of the lateral polarity proteins, *lgl* and *scrib*, doubles the frequency of germline invasion, as does removing both copies of Fasciclin II or RNAi knock down Neuroglian, both of which are the lateral adhesion factors (Bilder and Perrimon, 2000; Wei et al., 2004; Szafranski and Goode, 2006). By contrast, loss of one copy of *aPKC* or *crb* strongly suppresses invasion. Reducing the dosage of *gek* also decreases the frequency of invasion, consistent with a role for Gek in activating of NMYII laterally to stimulate the movement of the follicle cells into the germ line. Thus, these genetic interactions support the view that the invasive behaviour of *rhogap19d* mutant cells is driven by the expansion of the apical domain and Gek-dependent lateral contractility, both of which will increase the stress on the epithelium and promote buckling, without completely disrupting polarity.

**Figure 6.**
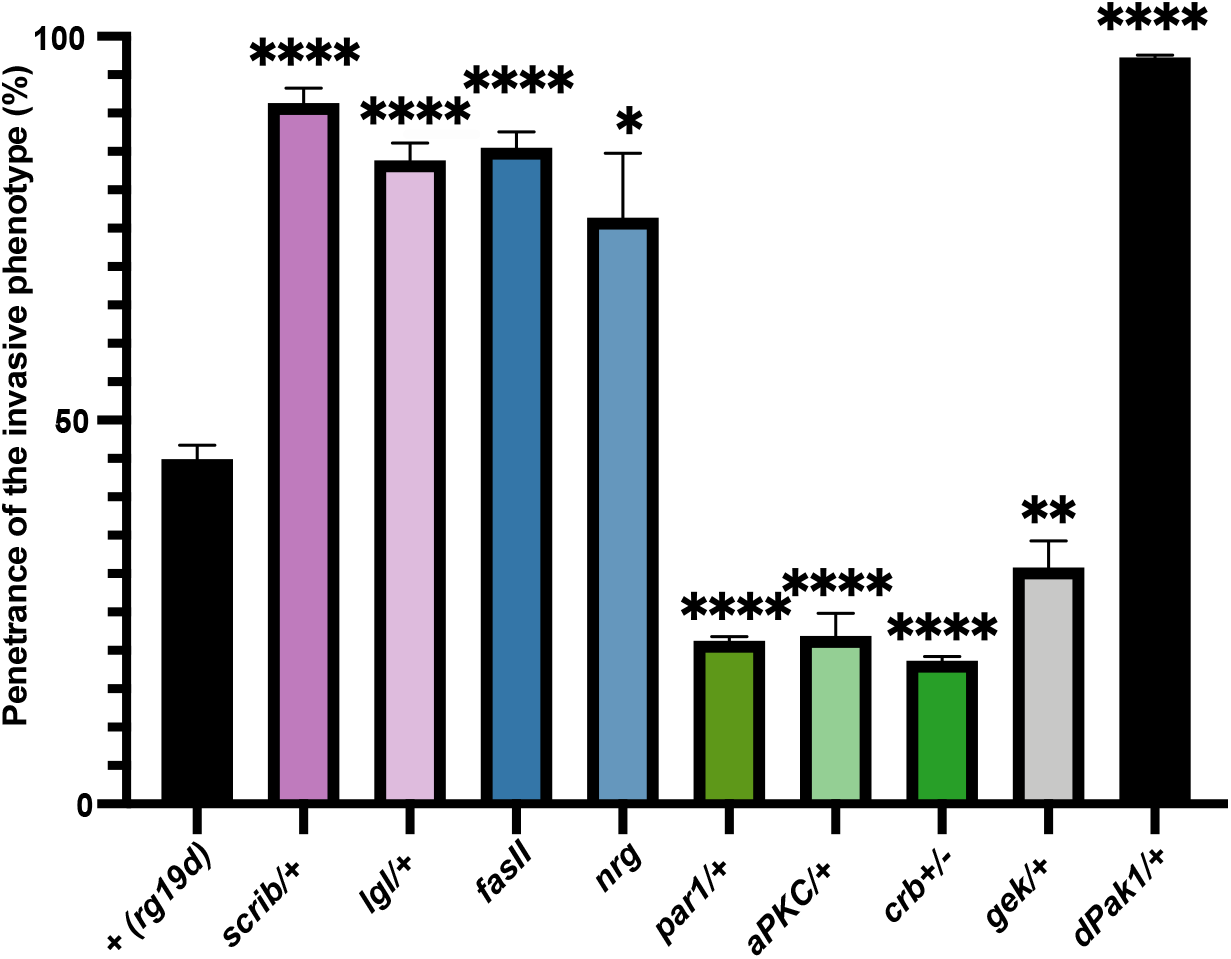
Genetic interactions between *rhogap19d* and other polarity factors. A histogram showing the penetrance of the germline invasion phenotype of large *rhogap19d* mutant clones in combination with other polarity mutants. Removing one copy of *scrib*, *lgl* or *Pak1* strongly enhances the penetrance of the invasion phenotype. *rhogap19d fasII* double mutant clones and *rhogap19d* clones in which *nrg* has been depleted by RNAi also show a highly penetrant invasive phenotype. Loss of one copy of *aPKC, gek*, *crb* or *par-1* strongly reduces the frequency of invasions.

Two of the mutants showed unexpected genetic interactions with *rhogap19d*. Firstly, reducing the dosage of the lateral polarity factor, Par-1, suppressed the invasive phenotype of the *rhogap19d* mutant, whereas the other lateral factors strongly enhance it. Par-1 localises to the lateral membrane and functions to limit the basal extent of the adherens junctions by phosphorylating and antagonising Bazooka (Par-3) (Benton and St Johnston, 2003; Wang et al., 2012). The ability of the *par-1* mutant to suppress *rhogap19d* indicates that Par-1 does not function in the same pathway as Scrib, Lgl, FasII and Nrg and suggests instead that it either negatively regulates these lateral factors or positively regulates apical ones. Secondly, Pak1 has been reported to function redundantly with aPKC to specify the apical domain downstream of active Cdc42 (Aguilar-Aragon et al., 2018). Although one would therefore expect the *pak1* mutant to suppress the invasive phenotype like mutants in the other in apical factors, it acts a strong enhancer of invasion. This is consistent with the role of Pak1 as a component of the lateral Scribbled complex and argues against the proposal that it functions as an apical Cdc42 effector kinase (Bahri et al., 2010).

## Discussion

Here we report that RhoGAP19D restricts Cdc42 activity to the apical side of the follicle cells and probably many other *Drosophila* epithelial tissues. In the absence of RhoGAP19D, both N-Wasp and Gek are recruited to the lateral membrane, indicating that Cdc42 is ectopically activated there. This implies that RhoGAP19D is the sole Cdc42GAP that represses Cdc42 laterally. This also suggests that the GEFs that activate Cdc42 are not restricted to the apical domain and can turn it on laterally once this repression is removed. This is consistent with the identification of multiple vertebrate GEFs with different localisations that contribute to apical Cdc42 activation (Oda et al., 2014; Otani et al., 2006; Qin et al., 2010; Rodriguez-Fraticelli et al., 2010; Zihni et al., 2014). Our results therefore identify RhoGAP19D as a new lateral polarity factor. This leads to a revised network of polarity protein interactions, in which RhoGAP19D functions as the third lateral factor that antagonizes the activity of apical factors, alongside Lgl, which inhibits aPKC, and Par-1 which excludes Bazooka/Par-3 (Wirtz-Peitz et al., 2008; Benton and St Johnston, 2003)(Fig 7a)

**Figure 7.**
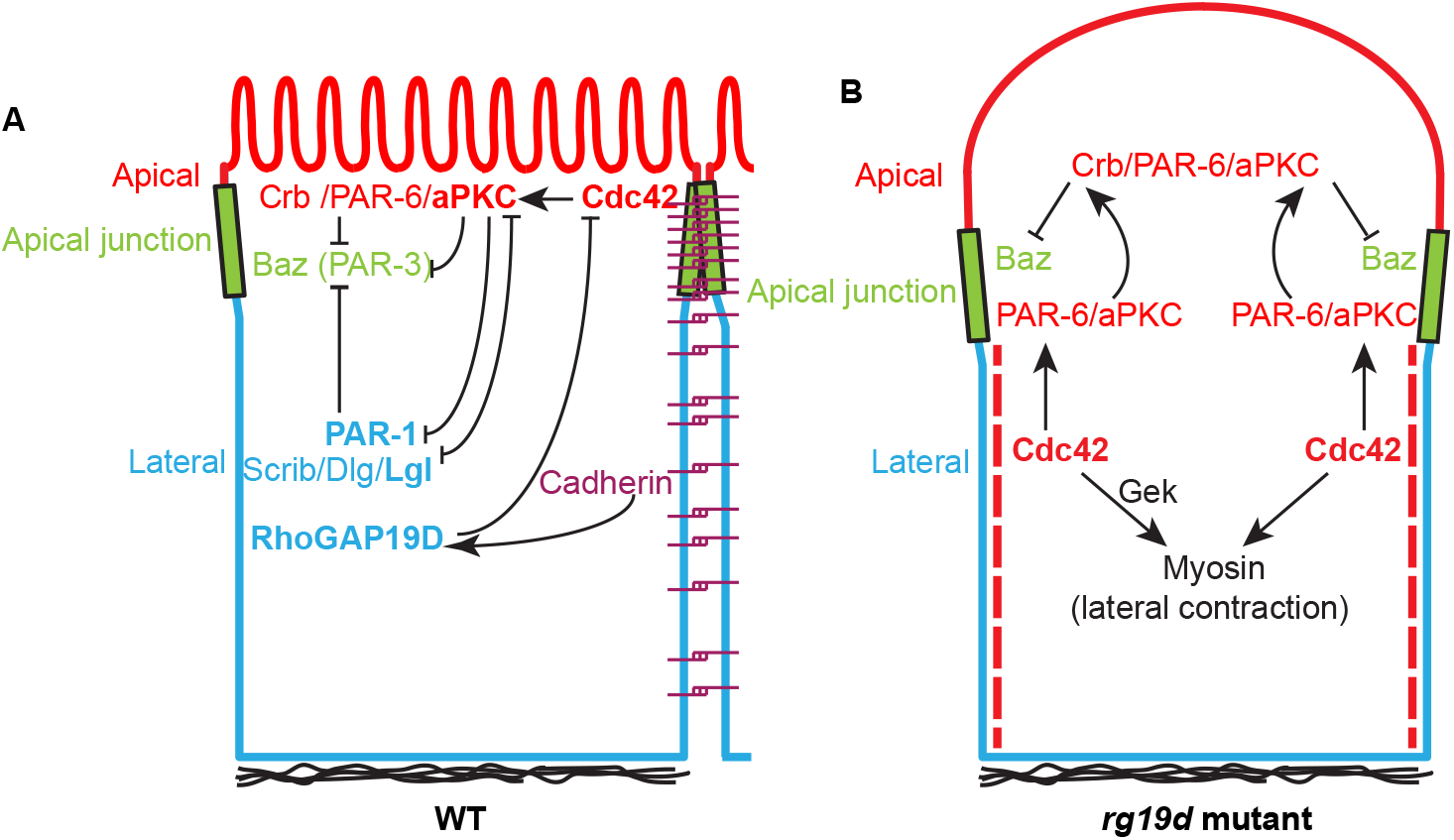
A revised network of inhibitory interactions between polarity factors. a) A diagram showing how the recruitment of RhoGAP19D to the lateral membrane by E-cadherin adhesion complexes restricts Cdc42 activity to the apical domain. This adds a new inhibitory interaction to the network of interactions between apical and lateral polarity factors. b) A model of the changes in *rhogap19d* mutant cells that lead to the invasive phenotype. In the absence of RhoGAP19D, active Cdc42 localises along the lateral, as well as the apical domain, and activates Gek to induce lateral actomyosin contractions. Lateral Cdc42-GTP also alters the conformation of Par-6/aPKC so that it is competent to bind to Crumbs. This primed Par-6/aPKC then diffuses until it binds to apical Crumbs, which activates aPKC’s kinase activity, resulting in expansion of the apical domain and a dome-shaped apical surface.

The function of RhoGAP19D is very similar to that of its orthologue PAC-1, which inhibits Cdc42 at sites of cell contact in early *C. elegans* blastomeres to generate distinct apical and basolateral domains (Anderson et al., 2008). Both RhoGAP19D and PAC-1 are recruited to the lateral domain by E-cadherin complexes, although the exact mechanism is slightly different. RhoGAP19D recruitment is strictly dependent on α-catenin, which links it through β-catenin to the E-cadherin cytoplasmic tail, whereas α-catenin (HMP-1) and p120-catenin (JAC-1) play partially redundant roles in recruiting PAC-1 to E-cadherin (HMR-1) in the worm (Klompstra et al., 2015).

Nevertheless, in both cases, the recruitment of the Cdc42GAP translates the spatial cue provided by the localization of cadherin to sites of cell-cell contact into a polarity signal that distinguishes the lateral from the apical domain. Classic work on the establishment of polarity MDCK cells grown in suspension has revealed that the recruitment of cadherin (uvomorulin) to sites of cell-cell contact is the primary cue that drives the segregation of apical proteins from basolateral proteins (Wang et al., 1990). Furthermore, the expression of E-cadherin in unpolarized mesenchymal cells is sufficient to induce this segregation, although the mechanisms behind this process are only partially understood (Wang et al., 1990; McNeill et al., 1990; Watabe et al., 1994; Nejsum and Nelson, 2007). Our observation that RhoGAP19D directly links cadherin adhesion to the polarity system in epithelial cells extends the results of Klompstra et al. (2015) in early blastomeres, strongly suggesting that PAC-1/RhoGAP19D plays an important role in the first steps in epithelial polarization.

Although PAC-1 and RhoGAP19D perform equivalent functions in early blastomeres and epithelial cells, there is one important difference between their mutant phenotypes. In *pac-1* mutants, Par-6 and aPKC are mislocalised to the contacting surfaces of *C. elegans* blastomeres where Cdc42 is ectopically active(Anderson et al., 2008). By contrast, Par-6 and aPKC are not mislocalised laterally in *rhogap19d* mutant *Drosophila* epithelial cells, even though lateral Cdc42-GTP does recruit two other Cdc42 effectors, N-Wasp and GEK. Thus, lateral Cdc42 activity is sufficient to recruit Par-6/aPKC to the lateral domain in early blastomeres, but not in epithelial cells. Instead, we observe that lateral Cdc42 activity in *rhogap19d* mutant follicle cells acts at a distance to expand the size of the apical domain. A likely explanation for this difference is the presence of Crumbs in epithelial cells. The interaction between Cdc42-GTP and Par-6 alters the conformation of Par-6 so that it can bind to Crumbs, which anchors the Par-6/aPKC complex to the apical membrane and activates aPKC’s kinase activity (Peterson et al., 2004; Whitney et al., 2016; Dong et al., 2020). Although Par-6 presumably binds to Cdc42 laterally in *rhogap19D* mutants and undergoes the conformational change, it cannot be anchored laterally in the absence of Crumbs. This activated Par-6/aPKC complex can then diffuse until it is captured by Crumbs in the apical domain, thereby increasing apical aPKC activity, providing an explanation for why the apical domain expands in *rhogap19d* mutant cells (Figure 7b). *C. elegans* has three Crumbs orthologues, but removal of all three simultaneously has no effect on viability or polarity (Waaijers et al., 2015). Thus, in contrast to epithelial cells, *C. elegans* Crumbs proteins are not required for Par-6/aPKC localisation and activation, suggesting that some other mechanism, such as Cdc42 binding is sufficient to activate aPKC.

If the failure of active Cdc42 to recruit aPKC laterally in *rhogap19d* mutant cells is due to the absence of Crumbs in this region, there must be a mechanism to exclude Crumbs from the lateral domain. One proposed mechanism depends on Yurt (Moe and EPB41L5 in vertebrates), which is restricted to the lateral domain by aPKC and binds to Crumbs to antagonize its activity (Laprise et al., 2006). However, we did not observe any lateral recruitment of aPKC in *rhogap19d; yurt* double mutant cells. Thus, there must be some parallel mechanism that excludes Crumbs, Par-6 and/or aPKC from the lateral domain.

Although loss of RhoGAP19D only leads to a partial disruption of polarity, it causes the follicular epithelium to invade the adjacent germline tissue with 40% penetrance. This invasive behaviour is not driven by an epithelial to mesenchymal transition, as the cells retain their apical adherens junctions and epithelial organisation. Instead, the deformation of the epithelium seems to be driven by the combination of an increase in lateral contractility and an expansion of the apical domain, as reducing the dosage of Gek, which activates myosin II to drive the contractility, significantly reduces the frequency of this phenotype, as does halving the dosage of any of the apical polarity factors. The invasion seems most likely to be driven by the buckling of the epithelium. Recent work has shown that epithelial monolayers under compressive stress and constrained by a rigid external scaffold have a tendency to buckle inwards (Trushko et al., 2020). The follicle cell layer is surrounded by an extracellular matrix that constrains the shape of the egg chamber and which should therefore resist expansion (Haigo and Bilder, 2011). Furthermore, the expansion of the apical domain should generate compressive stress because the domain is too long for the cells to adopt the lowest energy conformation while remaining in the epithelium. The lateral contractility may also contribute to compressive stress because reducing cell height while maintaining a constant volume will exert a pushing force on the neighbouring cells. In support of this view, lateral contractility has been shown to drive the folding of the imaginal wing disc between the prospective hinge region and the pouch (Sui et al., 2018). This phenotype therefore provides an example of how a partial disruption of polarity can induce cell shape changes that lead to major alterations in tissue morphogenesis (St Johnston and Sanson, 2011).

The *rhogap19d* phenotype resembles the defects earliest observed in the development of ductal carcinoma in situ (DCIS) (Halaoui et al., 2017). In flat epithelial atypia (FEA), the ductal cells are still organized into an epithelial layer, but they display apical protrusions that are strongly labeled by the apical polarity factor, Par-6. This suggests that the apical domain has expanded and bulges out of the cell, just as we observe in the *rhogap19d* mutant follicle cells. In the next stage, atypical ductal hyperplasia (ADH), the ductal cells start to invade the lumen of the duct, while retaining aspects of normal apical-basal polarity, again resembling the invasive phenotype reported here (Fig S4). Thus, these abnormalities, which can sometimes progress to DCIS and breast cancer, mirror the effects of lateral Cdc42 activation. The RhoGAP19D humans orthologues, ARHGAP21 and 23, has been shown to bind directly to α-catenin and localize to cell-cell junctions (Sousa et al., 2005) (Van Itallie et al., 2014). Furthermore, low expression of ARHGAP21 or 23 correlates with reduced survival rates in several cancers of epithelial origin (Fig S4) (Györffy et al., 2010). It would therefore be interesting to determine whether these orthologues perform the same functions in epithelial polarity as RhoGAP19D and if their loss contributes to tumour development.

## Materials & Methods

### Predicting Cdc42/GAP interactions

We identified and aligned putative *Drosophila* GAPs by searching the Pfam (Finn et al., 2016) hidden Markov model (HMM) profile PF00620.26 (RhoGAP) against the UniProt *Drosophila melanogaster* proteome using HMMsearch (Eddy, 2009). We aligned significantly scoring sequences together with human ARHGAP1 (from the 3D structure RCSB entry 1grn) using HMMalign (Eddy, 2009).

We scored the potential interaction of each *Drosophila* GAP/Cdc42 pair using the structure of human ARHGAP1/CDC42 (RCSB 1grn:B/A) via InterPReTS (Aloy & Russell, 2002), which assesses the effect of evolutionary changes at the interface structure using empirical pair potentials (Betts et al., 2015).

We modelled the Cdc42-RhoGAP19D 3D complex using SwissModel (Waterhouse et al., 2018) and rendered the interaction interface using the PyMOL Molecular Graphics System, Version 2.3.0, Schrödinger, LLC.

### Drosophila mutant stocks and transgenic lines

We used the following mutant alleles and transgenic constructs: *cdc42*^2^ ((Fehon et al., 1997); BDSC 9105), *p120*^ctn308^ (Myster et al., 2003); BDSC 81638), *shg*^IG29^ (BDSC 58471), *lgl*^4^ (Gateff, 1978); BDSC 36289), *scrib*^2^ (Bilder and Perrimon, 2000), *fasII*^G0336^ (Mao and Freeman, 2009); Kyoto 111871), *par-1*^6323^ (Shulman et al., 2000), *aPKC*^HC^ (Chen et al., 2018), *crb*^8F105^ (Tepass et al., 1990), *gek*^omb1080^(Gontang et al., 2011), *Pak1*^22^ (Newsome et al., 2000); a gift from Dickson lab), CadN^Δ14^ (Prakash et al., 2005), α-catenin RNAi (BDSC 33430), *nrg* RNAi (BDSC 38215), *fasII* RNAi (BDSC 34084), *fasIII* RNAi (BDSC 77396), *scrib* RNAi (BDSC 35748), *cora* RNAi (VDRC 9788), *rhogap19d* RNAi (P{TRiP.HMS00352}attP2; BDSC 32361), E-cad-EGFP (BDCS 60584; (Huang et al., 2009)), Lgl-EGFP (Tian and Deng, 2008), mNeonGreen-NWasp (a gift from Jenny Gallop, The Gurdon Institute), mCherryGrabFP-Baz (Harmansa et al., 2017), aPKC-EGFP (Chen et al., 2018), Zipper-EGFP (Lowe et al., 2014), UASp-GFP RhoGAP19D (BDSC 66167)., y^*^ w^*^; P{GawB}NP1624 (Traffic Jam-Gal4; (Brand and Perrimon, 1993)), nos-GAL4 (a gift from Ruth Lehmann). The following stocks were used to generate mitotic clones: ubiRFP-nls, hsflp, FRT20A4 (PBac{WH}f01417, Exelixis), FRT40A ubiRFP-nls (BDSC 34500), FRT82B, ubiRFP-nls (BDSC 30555) and FRT82B ubiGFP (BDSC 5188), y w hs-FLP;; Act5C>CD2>Gal4, UAS:mRFPnls ((Pignoni and Zipursky, 1997); BDSC 30558).

#### Generation of endogenously tagged RhoGAP19d

The mNeonGreen tag (Shaner et al., 2013) was fused to the N-terminus of RhoGAP19d by CRISPR-mediated homologous recombination. In vitro synthesised gRNA to a CRISPR target (target sequence GGTGGCGACTCCGGCAGCGGCGG, CRISPR; loc. 25985bp from the 5’ end) and a plasmid donor containing the ORF of mNeonGreen as well as appropriate homology arms (1.5 kb upstream and downstream) were co-injected into nos-Cas9–expressing embryos (BDSC 54591). Single F0 flies were mated to y w flies and allowed to produce larvae before the parents were analysed by PCR. Progeny from F0 flies in which a recombination event occurred (as verified by PCR) were crossed and sequenced to confirm correct integration. Several independent mNeonGreen-RhoGAP19d lines were isolated. Recombinants carry the mNeonGreen coding sequence inserted immediately downstream of the endogenous start codon with a short linker (Gly-Ser-Gly-Ser) between the coding sequence of mNeonGreen and the coding sequence of RhoGAP19d. Homozygous flies are viable and healthy.

To discriminate between the signal of mNeonGreen-RhoGAP19D in the germline and in the somatic follicle cells, UAS-RhoGAP19D RNAi was expressed in the germline under the control of nos-Gal4. The knock-down was efficient, which allowed visualisation Neon-RhGAP19D expression in the follicle cells only.

#### Generation of *rhogap19d* mutant flies

We used the error prone repair of CRISPR/Cas9-induced dsDNA breaks by the nonhomologous end joining repair pathway to generate null alleles of *rhogap19d* (Bassett et al., 2013). The following target sequence was used GGGTCGGGATCCCTTTCGGGGGG, CRISPR (38613bp from 5’ end). *In vitro* synthetized gRNA to the target sequence and Cas9 mRNA were injected into FRT20A4 embryos. Single F0 flies were mated to y w flies. DNA from F0 flies was extracted and analysed by High Resolution Melting (HRM). Short fragments of about 150bp covering the region containing the target sequence were amplified by PCR. Slow melting curves were generated for the PCR products and changes in sequence were measured by changes in fluorescence as the strands separate. This technique allows the detection of single base changes. Progeny of promising F0 candidates were balanced, and sequenced. Multiple mutant alleles of *rhogap19d* on the FRT20A4 chromosome were isolated and analysed. The mutants that contain insertions or deletions generating premature STOP codons were kept for clonal analysis. These mutations generate proteins of ~436 amino acids that lack the RhoGAP domain, but contain the PDZ domain. *rhogap19d* mutant flies are semi-lethal. The follicle cell phenotypes of the *rhogap19d* alleles used in this study are rescued by the expression of UAS-RhoGAP19D-GFP under the control of TJ-Gal4, confirming that they are caused by loss of RhoGAP19D function.

### Reagents

The following antibodies were used: Primary antibodies: anti-Armadillo (N2 7A1, Developmental Studies Hybridoma Bank (DSHB) 1:100 dilution), anti-aPKC (C-20, sc-216-G, Santa Cruz, goat polyclonal IgG, 1:500), anti-Cno ((Takahashi et al., 1998); a gift from M. Peifer, University of North Carolina, USA, 1:1,000 dilution) anti-Gek ((Gontang et al., 2011); a gift from the Clandinin lab,1:25 dilution), anti-PH3 (Cell Signalling #9701S, 1:500 dilution).

Secondary antibodies: Alexa Fluor secondary antibodies (Invitrogen) were used at a dilution of 1:1,000. Alexa Fluor 488 goat anti-mouse (#A11029), Alexa Fluor 488 goat anti-rabbit (#A11034), Alexa Fluor 647 goat anti-mouse (#A21236), Alexa Fluor 647 goat anti-rabbit (#A21245).

F-Actin was stained with phalloidin conjugated to Rhodamine (Invitrogen, Cat. #R415, 1:500 dilution). The cell membranes were labelled with CellMask Orange Plasma Membrane Stain or CellMask Deep Red Plasma Membrane Stain (Thermo Fisher Scientific).

### Immunostaining

Ovaries from fattened adult females, salivary glands from L3 instar larvae and accessory glands from virgin males were dissected in Phosphate-buffered Saline (PBS) and fixed with rotation for 20 min in 4% paraformaldehyde and 0.2% Tween 20 in PBS. After a few washes with PBS-0.2%Tween, tissues were then incubated in 10% bovine serum albumin (BSA) in PBS to block for at least 1 hour at room temperature. Incubations with primary antibodies were performed at 4°C overnight in PBS, 0.2% Tween 20 and 1% BSA. This step was followed by four washes with PBS- 0.2% Tween and samples were then incubated for 3-4 h with secondary antibody at room temperature. Specimens were then washed several times in washing buffer and mounted in Vectashield containing DAPI (Vector Laboratories).

For stainings with the anti-Gek antibody, ovaries were heat fixed as described by (Chen et al., 2018).

### Imaging

Fixed samples and live imaging was performed using an Olympus IX81 (40×/1.3UPlan FLN Oil or 60×/1.35 UPlanSApo Oil) or a Leica SP8 white Laser (63×/1.4 HC PL Apo CS Oil) inverted confocal microscope. For live observations, ovaries were dissected and imaged in 10S Voltalef oil (VWR Chemicals). Images were processed with Fiji (Schindelin et al., 2012) or Leica analysis software.

#### *Drosophila* genetics

Standard procedures were used for Drosophila maintenance and experiments. Flies were grown on standard fly food supplemented with live yeast at 25 °C. Follicle cell clones were induced by incubating larvae or pupae at 37°C for 2 hours every 12 hours over a period of at least 3 days. Adult females were dissected at least 2 days after the last heat shock. In some experiments adult flies were heat-shocked for at least 3 days and dissected one day after the last heat shock.

We used the Flipout technique with Actin5c>Cd2>Gal4 to generate marked clones of cell expressing RNAi constructs (Fig. S3). Flp recombination was induced by incubating larvae or pupae at 37°C for 2 hours every 12 hours over a period of at least 3 days.

### Genetic interactions

To test for genetic interactions between *rhogap19d* and adhesion molecules or polarity factors, we analysed the frequency of follicle cell invasions at stages 7 and 8 in large anterior *rhogap19d* clones that covered at least 25% of the follicular epithelium in each genetic background.

An unpaired, two-tailed students t-test with Welch’s correction was used to determine whether any differences between the penetrance of the invasion phenotype in *rhogap19d* alone and in combination with each mutant or RNAi knockdown were significant.

### Quantifications of the total number of follicle cells per egg chamber in *rhogap19d* mutants

Confocal z-stacks of whole egg chambers were collected on a Leica SP8 white laser microscope. Each egg chamber was divided in three regions. Nuclei were counted twice per region.

### Reproducibility of experiments

All experiments were repeated multiple times as listed below. For each figure, the first number indicates the number of times that the experiment was repeated and the second indicates the number of egg chambers or clones analysed. The number of independent experiments performed were: Figure 1a (3; 28), Figure 2c (5; 89), Figure 2e (3; 34), Figure 2f (2; 20), Figure 2h (7; 56), Figure 2i (4; 34), Figure 2j&k (3; 6), Figure 2l (2; 9), Figure 2m (3; 12), Figure 2n (2; 7). Figure 3a (4; 24), Figure 3b (3; 46). Figure 4a (7; 67), Figure 4b (3; 78), Figure 4c (5; 98), Figure 4d (2; 16), Figure 4e (3; 56). Figure 5a (7, 56), Figure 5c (5; 47), Figure 5d (3; 98), Figure 5e (3; 18). Figure 6: *rhogap19d* (8; 301), *scrib/+* (3; 194), *lgl/+* (3; 94), *fas2* (3; 185), *nrg* (3; 114), *par1/+* (3; 155), *aPKC/+* (4; 98), *crb* (2; 129), *gek* (2; 78), *Pak1* (3; 228).

**Table 1.** List of GAPs in *Drosophila*. The table shows all known GAPs with their additional names and UniProt numbers.

| Gene symbol | Gene name | Other names | UniProt |
| --- | --- | --- | --- |
| CdGAPr | protein-related | GAP, d-CdGAPr | Q9VIS1 |
| conu | conundrum |  | Q8T0G4 |
| cv-c | crossveinless c | RhoGAP88C | A8JR05 |
| Graf | GRAF ortholog (h. sapiens) |  | X2JDY8 |
| Ocrl | syndrome of Lowe | EG:86E4.5 | Q95R90 |
| RacGAP84C | protein at 84C | rnRacGAP | P40809 |
| RhoGAP1A | protein at 1A | EG:23E12.2 | Q6W436 |
| RhoGAP5A | protein at 5A |  | Q9W4A9 |
| RhoGAP15B | protein at 15B |  | Q0KHR5 |
| RhoGAP16F | protein at 16F |  | Q9VWY8 |
| RhoGAP18B | protein at 18B | whir | Q9VWL7 |
| RhoGAP19D | protein at 19D |  | Q9VRA6 |
| RhoGAP54D | protein at 54D |  | A1ZAW3 |
| RhoGAP68F | protein at 68F | CG 6811 | M9PC96 |
| RhoGAP71E | protein at 71E | l(3)j6B9 | B7Z058 |
| RhoGAP92B | protein at 92B |  | A0A0B4LHC1 |
| RhoGAP93B | protein at 93B | CrGAP | Q9VDE9 |
| RhoGAP100F | protein at 100F | Syd-1 | Q9V9S7 |
| RhoGAP102A | protein at 102A | Dm_4:1183 | H9XVN1 |
| RhoGAPp190 | protein at 190 | p190RhoGAP, p190 RhoGAP | Q9VX32 |
| Rlip | Ral interacting protein | dRalBP, D-RLIP | Q9VDG2 |
| tum | tumbleweed | racGAP50C, acGAP, RacGAP, DRacGAP | Q9N9Z9 |

**Table 2.** GAPs ranked by the predicted strength of their interaction with Cdc42. RhoGAP19D has the highest Z-score.

| **Protein 1** | **Gene 1** | **%id1** | **Protein 2** | **Gene 2** | **%id2** | **PDB** | **Z-score** |
| --- | --- | --- | --- | --- | --- | --- | --- |
| Q9VRA6-RhoGAP | RhoGAP19D | 24 | P40793-RAS | Cdc42 | 94 | 1AM4 | 3.329 |
| M9PC96-RhoGAP | RhoGAP68F | 40 | P40793-RAS | Cdc42 | 92 | 1GRN | 3.122 |
| Q9VIS1-RhoGAP | CdGAPr | 26 | P40793-RAS | Cdc42 | 92 | 1GRN | 2.999 |
| A0A0B4LHC1-RhoGAP | RhoGAP92B | 36 | P40793-RAS | Cdc42 | 92 | 2NGR | 2.9 |
| P40809-RhoGAP | RacGAP84C | 26 | P40793-RAS | Cdc42 | 92 | 1GRN | 2.88 |
| Q8T0G4-RhoGAP | conu | 26 | P40793-RAS | Cdc42 | 92 | 1GRN | 2.835 |
| Q9VDE9-RhoGAP | RhoGAP93B | 30 | P40793-RAS | Cdc42 | 92 | 1GRN | 2.823 |
| X2JDY8-RhoGAP | Graf | 29 | P40793-RAS | Cdc42 | 94 | 1AM4 | 2.748 |
| Q0KHR5-RhoGAP | RhoGAP15B | 28 | P40793-RAS | Cdc42 | 94 | 1AM4 | 2.723 |
| A8JR05-RhoGAP | cv-c | 29 | P40793-RAS | Cdc42 | 94 | 1AM4 | 2.642 |
| Q9VWL7-RhoGAP | RhoGAP18B | 28 | P40793-RAS | Cdc42 | 94 | 1AM4 | 2.488 |
| Q9VWY8-RhoGAP | RhoGAP16F | 22 | P40793-RAS | Cdc42 | 94 | 1AM4 | 2.142 |
| Q9N9Z9-RhoGAP | tum | 22 | P40793-RAS | Cdc42 | 94 | 1AM4 | 1.951 |
| Q9VX32-RhoGAP | RhoGAPp190 | 28 | P40793-RAS | Cdc42 | 92 | 1GRN | 1.93 |
| Q9VDG2-RhoGAP | Rlip | 32 | P40793-RAS | Cdc42 | 92 | 2NGR | 1.683 |

## Supplemental Information

**Table S1**: List of CRISPR-mediated mutations in candidate Cdc42 GAPS

**Figure S1**:RhoGAP92B, RhoGAP68F, CdGAPr, RacGAP84C, Conu and RhoGAP93B are not required for follicle cell polarity.

**Figure S2**: RhoGAP19D localises laterally in multiple epithelia.

**Figure S3**: Mutants that do not affect the localisation of RhoGAP19D

**Figure S4**: The *rhogap19d* phenotype resembles the early steps in breast cancer.

**Movie: S1**: A time-lapse movie of a stage 7 egg chamber containing a large *rhogap19d* mutant clone (marked by the loss of RFP; magenta) and expressing GFP-aPKC (green). Frames were captured every 15s. Elapsed time, 11 mins.

## Acknowledgements

We thank Jenny Gallop, Franck Pichaud, Barry Dickson, Marc Peifer, Thomas Clandinin, Ruth Lehmann, Andrea Brand and their labs for fly stocks and for antibodies; the Developmental Studies Hybridoma Bank (DSHB), the Kyoto Stock Center (DGRC), the Vienna Drosophila Resource Center (VDRC), Exelixis, and the Bloomington *Drosophila* Stock Center (BDSC), Santa Cruz Biotechnology for antibodies and fly stocks; Jia Chen for help with heat fixation, Richard Butler for help with image analysis, Nick Lowe for help with HRM analysis and members of the D. St J. lab for technical assistance, helpful comments and criticism. RBR acknowledges support from the German Network of Bioinformatics Infrastructure (de.NBI) funded by the Bundesministerium für Bildung und Forschung (BMBF).

## Author contributions

The project was conceived by W.F., R.B., R.B.R. and D. St J. The structural modelling of *Drosophila* RhoGAP proteins was performed by F.R and R.B.R. The *Drosophila* experiments were performed by W.F, R.B. and E.L. and the data were analysed by W.F, R.B. and D. St J. The mNeonGreenWASP line was generated by Y.I. and J.G. The project funding, administration and supervision were provided by D. St J. and R.B.R. W.F and D. St J. prepared the figures and wrote the manuscript, which was edited and reviewed by all authors.

