## Supplemental info for "RhoGAP19D inhibits Cdc42 laterally to control epithelial cell shape and prevent invasion" (from another version of this preprint: version 1, 2020-09-17)

**Table S1**

| Gene | Expression | Viability | Mutation | Predicted protein |
|---|---|---|---|---|
| RhoGAP19D | moderate | semi-lethal | 1) 5nt deletion and 4nt insertion | 436aa, missing GAP domain |
| | | | 2) 7nt deletion | 448aa, missing GAP domain |
| | | | 3) 13nt deletion | 446aa, missing GAP domain |
| RhoGAP68F | high | viable | 4nt deletion | 149 aa, missing all domains |
| CdGAPr | moderate | semi-lethal | 5nt deletion and 6nt insertion | 158 aa, missing all domains |
| RhoGAP92B | moderately high | lethal | 2 nt deletion | 329 aa, missing GAP domain |
| RacGAP84C | extremely low | viable | 35 nt deletion | 79 aa, missing all domains |
| Conu | moderately high | viable | 4 nt deletion and 11 nt insertion | 98 aa, missing all domains |
| RhoGAP93B | moderate | viable | 5nt deletion and 6nt insertion | 53 aa, missing all domains |

Table 3. Characterisation of mutants generated by CRISPR.

### Figure S1

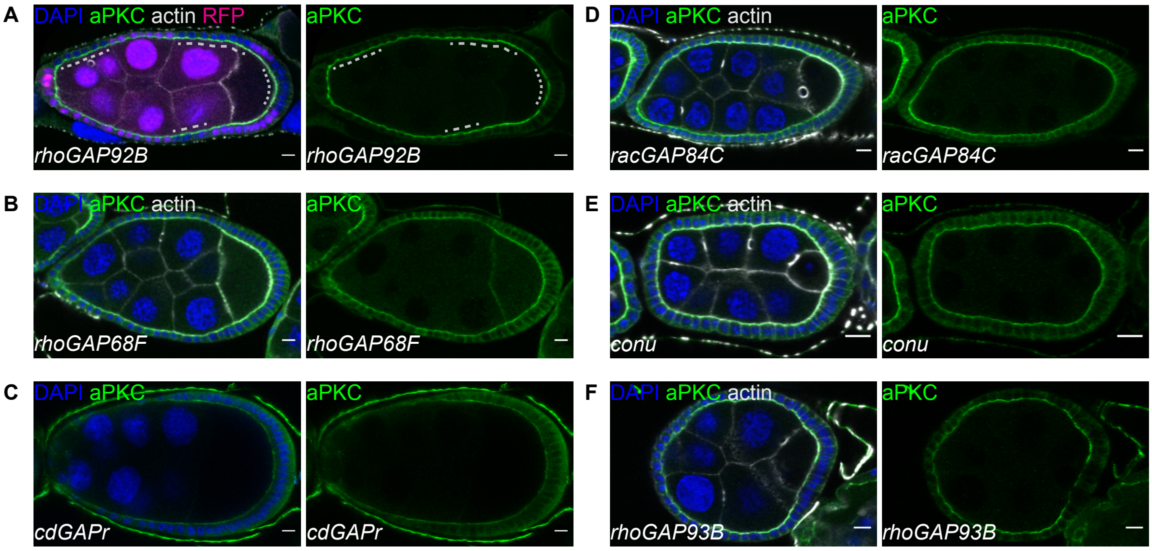


**Fig. S1. RhoGAP92B, RhoGAP68F, CdGAPr, RacGAP84C, Conu and RhoGAP93B are not required for follicle cell polarity.** (A) Egg chambers containing *rhoGAP92B* mutant cells marked by loss of RFP. Loss of RhoGAP92B does not affect aPKC localisation (A; *n*=28) or Lgl localisation (not shown; *n*=24). Dashed lines indicate RFP-negative mutant cells. (B-F) Egg chambers from flies homozygous for *rhoGAP68F* (B; *n*=109), *cdGAPr* (C; *n*=12), *racGAP84C* (D; *n*=104), *conu* (E; *n*=90) and *rhoGAP93B* (F; *n*=119) mutant show normal localisation of aPKC (B-F) and Lgl (not shown) and normal organisation of the follicular epithelium. Scale bars: 10 µm.

### Figure S2

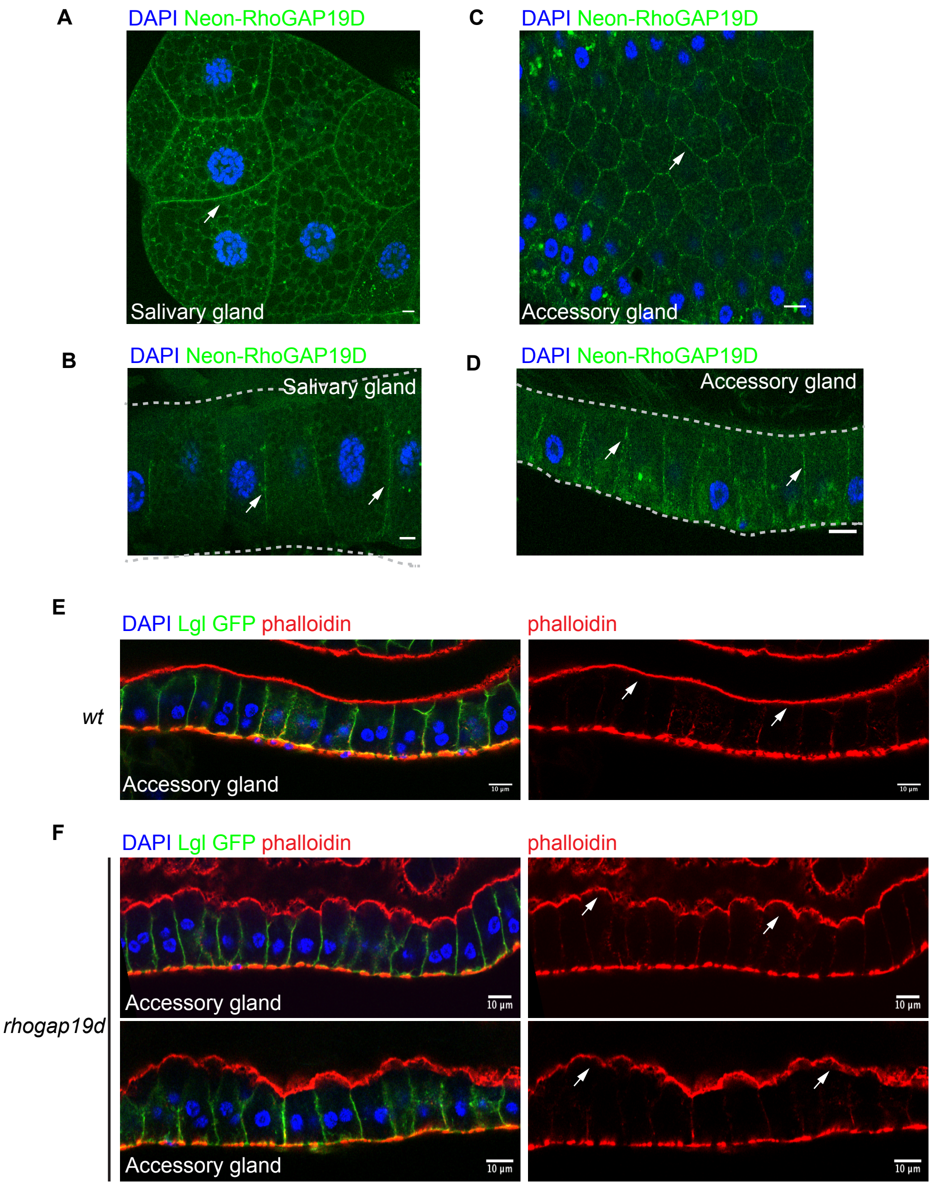


**Fig. S2 RhoGAP19D localises laterally in multiple epithelia.**

a) A top view and a cross section (b) through a salivary gland from an L3 larva, showing mNeonGreen-RhoGAP19D (green) localisation to the lateral domain. DAPI (blue), n=8 larvae.
c) A top view and a cross section (d) through an accessory gland from a mated male. mNeonGreen-RhoGAP19D (green) localises laterally. DAPI (blue), n=10 males.
e) Cross section through an accessory gland of a vigin wild-type male, Lgl-GFP (green), phalloidin (red), DAPI (blue). Arrows indicate the apical side of the epithelium, n=6 males.
f) Two examples of an accessory gland from virgin *rhogap19d* mutant males, Lgl-GFP (green), phalloidin (red), DAPI (blue). The apical side (white arrows) protrudes into a dome and the cells are much taller than in the wild-type tissue. n=11 males. Scale bars 10μm.

**Figure S3**

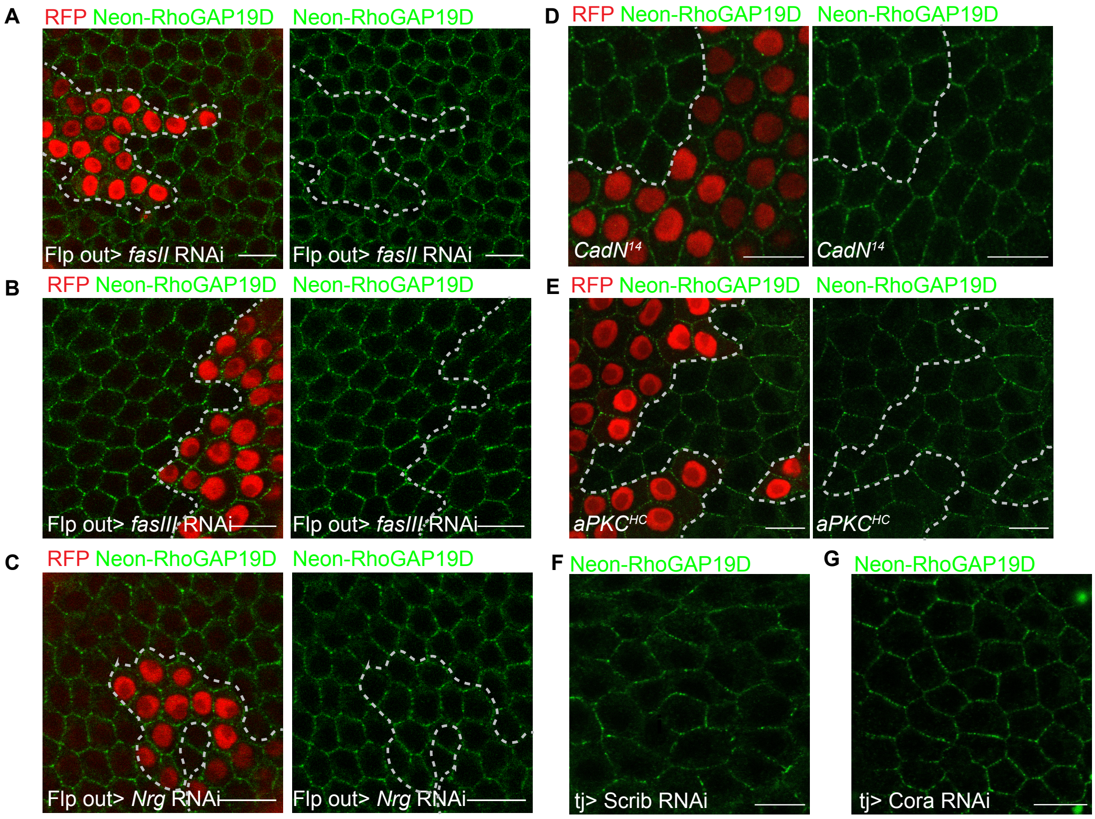


**Figure S3 Mutants that do not affect the localisation of RhoGAP19D**

a-c) Wild-type localisation of mNeonGreen-RhoGAP19D at the lateral cortex of stage 7 follicle cells in which *fasII* (a; n=21), *fasIII* (b; n=19) and *nrg*(c; n=25) have been knocked down by UAS-RNAi Flp out system (cells that express RFP coexpress RNAi constructs).

Each experiment was performed three independent times.

d) A stage 8 egg chamber with a clone of cells mutant for *N-cadherin1* and *N-cadherin2* (marked by the loss of RFP). mNeonGreen-RhoGAP19D localises normally in the mutant cells. (n=15, Experiment performed twice).

e) mNeonGreen-RhoGAP19D localises to the cortex in *aPKC*$^{HC}$ clones at stage 8 (marked by the loss of RFP). n=16, experiment performed twice.

f) mNeonGreen-RhoGAP19D is correctly localised in cells treated with RNAi against *scrib* n=23 or *coracle* (g) n=16. Stage 8. Experiments performed twice.

Scale bars 10$\mu$m.

### Figure S4

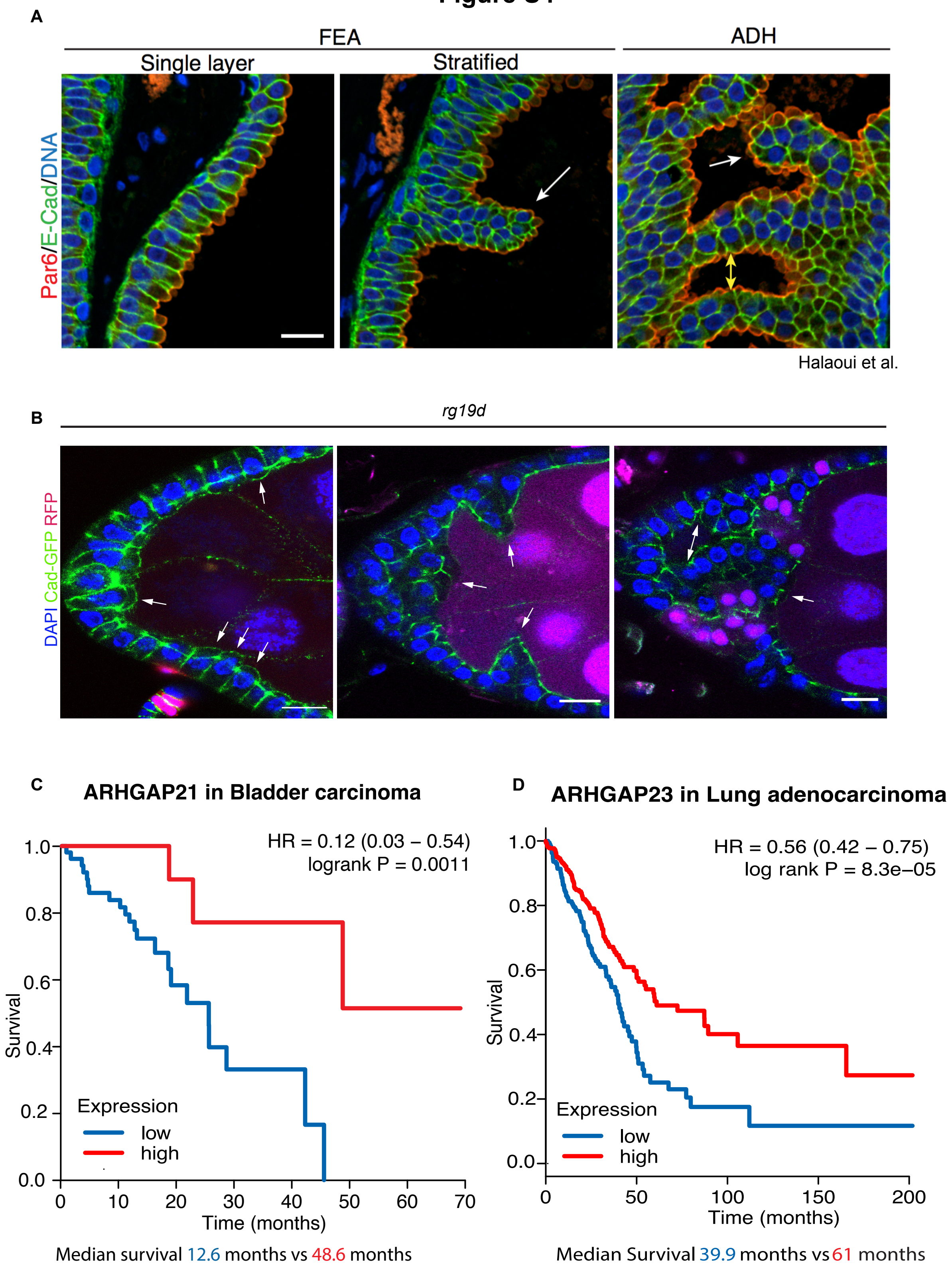


**Figure S4 The *rhogap19d* phenotype resembles the early steps in breast cancer.**
a) Images of breast tissue samples reproduced from Halaoui et al. (2017) Supplementary Figure 3, showing examples of Flat Epithelia Atypia (FEA) and Atypical Ductal Hyperplasia (ADH). Samples were immuno-stained for Par6 (red), E-cad (green) and DAPI (blue).
b) *rhogap19d* mutant cells (marked by the absence of RFP), stained for E-cad-GFP (green) and DAPI (blue), show similar apical bulges and invasions to FEA and ADH. Cells first bulge apically (white arrows in the left panel), then start to collectively invade the germline (middle panel), to finally form big clusters inside the egg chamber (right panel). Stage 8 egg chamber. Experiment repeated 5 times, n=68. Scale bars 10μm.
c) Kaplan-Meier survival plot for ARHGAP21 expression (high; top quartile vs low; bottom quartile) in bladder carcinoma. Survival data was retrieved from the kmplot resource (kmplot.com) described in Gyöffrey et al. (2010)
d) Kaplan-Meier survival plot for ARHGAP23 expression in lung adenocarcinoma.
